## Supplementary Information for "Comparison of state-of-the-art error-correction coding for sequence-based DNA data storage"

### Supplementary Notes

#### Supplementary Note 1: Patches to codec implementations

In order to ensure representative error-correction performance or facilitate automated testing, the implementations of the DNA-Aeon, Goldman, HEDGES, and Yin-Yang codecs were patched. The patches are available in the GitHub repository at github.com/fml-ethz/dt4dds-benchmark, and outlined in the following.

##### DNA-Aeon

For DNA-Aeon, a custom codebook was used to change homopolymer constraints. In addition, the encoding script was adjusted to enable a local python runtime environment, as well as support custom paths for in- and output files. Moreover, intermediate files generated during encoding were renamed to prevent race conditions due to name clashes.

##### Goldman

The encoding pipeline was modified to record the total number of segments used for encoding. This information was then used in the decoding pipeline to filter out invalid segments. Moreover, the decoding pipeline was modified to support arbitrary binary files, filter out corrupted segments, as well as join overlapping segments to yield consensus sequences by majority voting prior to segment decoding.

##### HEDGES

The implementation at the official GitHub repository[1] has been changed to a pure C++ implementation in November 2024, from the original implementation as C++ modules callable from Python 2. Unfortunately, the old implementation has been wiped from the repository during the re-implementation. Therefore, a fork of the old implementation must now be used, such as github.com/shulp2211/hedges. This study is using the old, original implementation throughout. Individual en- and decoding scripts were set up, following the pipelines used in the provided test script. In addition, these scripts were changed to support arbitrary file sizes by padding the last packet with sentinel values. As a result, effective code rate depends on input file size, due to varying completeness of the last packet. As the decoding depends on the presence of specific adapter sequences up- and downstream of the data-carrying nucleotides, these are added and removed to all sequences during de- and encoding respectively.

##### Yin-Yang

A script file for implementing the en- and decoding tasks described in the README was set up to serve as the entry point for their automation. In addition, supplementary files required for decoding were saved separately after encoding and automatically re-supplied to the script during the decoding step.

#### Supplementary Note 2: Considerations for reporting storage density

We chose to report code rates and storage densities without the overhead of amplification adapters or other sequence components unrelated to the data encoding in this study. This means both metrics only consider the codec's sequence output incl. indexes, redundancy, etc. as the nucleotide count used for encoding. Besides simplicity, the main reasons for this were the absence of any sequence overhead in the synthetic benchmarks, as well as the low relevance to the comparison between codecs in this study (e.g., codecs had identical amplification adapters). Further complicating the matter, our *in-vitro* replication required padding the output of some codecs to achieve a homogeneous sequence length (see Supplementary Fig. 8). As experimental considerations rather than codec constraints necessitated this padding, including it in the calculations for storage density would have unduly affected codecs which produced shorter sequences.

A major downside of this choice is the comparableness outside of the codecs in this study. While this is a common problem in the DNA data storage literature due to differences in the choice of amplification adapters or the support for random access, it impedes fair comparisons. Thus, we also report our experimental storage densities and those from the literature using other definitions of the nucleotide count in Supplementary Table 9. In all cases, a molecular weight of 662 g $mol^{-1}$ $bp^{-1}$ (i.e., double-stranded DNA at 616 g $mol^{-1}$ $bp^{-1}$ with two sodium counterions at 2x23 g $mol^{-1}$ $bp^{-1}$) is assumed, leading to the following equations:

$$\text{Code rate in bit nt}^{-1} = \frac{\text{File size in bit}}{\text{Nucleotide count}}$$

$$\text{Storage density in EB g}^{-1} = \frac{\text{Code rate in bit nt}^{-1} \times \frac{1}{8}\text{ byte bit}^{-1} \times 10^{-18}\text{ EB byte}^{-1}}{662\text{ g mol}^{-1}\text{ bp}^{-1} \times N_A^{-1} \times 1\text{ bp nt}^{-1} \times \text{Physical redundancy}}$$

This simplifies to:

$$\text{Storage density in EB g}^{-1} = \frac{\text{Code rate in bit nt}^{-1}}{\text{Physical redundancy}} \times 113.7\text{ EB bit}^{-1}\text{ nt g}^{-1}$$

Note that the exact molecular weight used for calculations varies slightly between studies (e.g., Organick et al.[2] use 325 g $mol^{-1}$ $nt^{-1}$ with single-stranded DNA and assume $1024^6$ byte $EB^{-1}$). To harmonize the results shown in Supplementary Table 9, results from literature were re-calculated using the above definitions (i.e., using 331 g $mol^{-1}$ $nt^{-1}$ for single-stranded DNA and $1000^6$ byte $EB^{-1}$). This leads to slight differences in the listed storage densities compared to those reported in the original studies.

#### Supplementary Note 3: Variations in sequencing depth and coverage

In the experimental replications of the low- and high-fidelity workflows, it was assumed that each codec's sequences were homogeneously represented in the oligo pools, if they were synthesized simultaneously. To ensure this, the order of the sequences supplied to the commercial synthesis companies was randomized, thereby precluding any chip-related bias.[3] Nonetheless, the sequencing data strongly suggests the presence of a systematic bias between codecs, highlighted by the inhomogeneity of sequencing depths (see Supplementary Fig. 9).

As outlined in the main manuscript, it is inconclusive which process (i.e., synthesis, amplification, or sequencing) specifically caused this bias. However, given the randomization during synthesis (see above) and the presence of this bias in both synthesis technologies, it is unlikely to be related to synthesis. PCR is known to have biased amplification,[3–6] which would explain the difference in the bias's severity between the low- and high-fidelity scenario (due to different numbers of PCR cycles in the two workflows).

Plausible causes for the PCR-induced bias are sequence features introduced by codecs (e.g., repetitive elements in the Goldman codec, indexing regions) or the barcodes added to each codecs' sequences (see Supplementary Fig. 8 and Supplementary Table 10). More experiments would be needed to conclusively elucidate the origin of the inhomogeneity. However, the observed bias correlates strongly with the rate of sequence loss in the sequencing data (see Supplementary Fig. 9), thereby likely causing the systematic deviations between simulated predictions and the experimental results. Specifically, amplification after synthesis likely caused specific enrichment of sequences from some codecs in the oligonucleotide pools (see Sequencing depth of 1000x samples in Supplementary Fig. 9). Then, during dilution, enriched sequences were more likely to be sampled. This results in a lower probability for sequence dropout during sequencing, reducing the need for logical redundancy for the decoder.

### Supplementary Tables

**Supplementary Table 1: Full results of clustering performance.** For each clustering algorithm, multiple parameter sets were tested, if supported. These parameter sets deviated from the default settings in the parameters described in the Parameters column. Each parameter set was tested once with experimental data from electrochemical synthesis (Elec.), and once with experimental data from synthesis by material deposition (M.D.), see Methods for additional details and definitions of sensitivity, similarity, and specificity. The best-performing parameter set of each clustering algorithm is highlighted with green shading. Note that MMseqs2 was not suited for further consideration and pairing with codecs due to common violation of the memory constraint.

| Clustering | Parameters | Scenario | Sensitivity | Similarity | Specificity | Time / s |
|---|---|---|---|---|---|---|
| Naïve | None | Elec. | 1.000 | 0.997 | 0.057 | 3 |
| | | M.D. | 1.000 | 1.000 | 0.084 | 2 |
| CD-Hit | Default | Elec. | 1.000 | 0.999 | 0.334 | 161 |
| | | M.D. | 1.000 | 1.000 | 0.895 | 145 |
| | Identity threshold 80%<br>Word size 5 | Elec. | 1.000 | 0.999 | 0.959 | 4152 |
| | | M.D. | 1.000 | 1.000 | 0.996 | 3859 |
| | Identity threshold 85%<br>Word size 6 | Elec. | 1.000 | 0.999 | 0.893 | 278 |
| | | M.D. | 1.000 | 1.000 | 0.990 | 243 |
| Clover | Depth 10 | Elec. | 0.381 | 0.953 | 0.611 | 207 |
| | | M.D. | 0.295 | 0.965 | 0.618 | 181 |
| | Depth 15<br>Horizontal drift 5 | Elec. | 0.987 | 0.998 | 0.656 | 195 |
| | | M.D. | 0.981 | 1.000 | 0.992 | 153 |
| | Depth 15<br>Vertical drift 4 | Elec. | 0.988 | 0.999 | 0.657 | 179 |
| | | M.D. | 0.987 | 1.000 | 0.994 | 151 |
| | Depth 20 | Elec. | 0.990 | 0.999 | 0.594 | 192 |
| | | M.D. | 0.998 | 1.000 | 0.993 | 145 |
| | Depth 20<br>Horizontal drift 5 | Elec. | 0.990 | 0.999 | 0.594 | 217 |
| | | M.D. | 0.998 | 1.000 | 0.993 | 149 |
| | Depth 20<br>Vertical drift 4 | Elec. | 0.990 | 0.999 | 0.593 | 203 |
| | | M.D. | 0.997 | 1.000 | 0.993 | 152 |
| | Default | Elec. | 0.988 | 0.999 | 0.654 | 189 |
| | | M.D. | 0.989 | 1.000 | 0.994 | 144 |
| LSH | Default | Elec. | 0.987 | 0.999 | 0.190 | 770 |
| | | M.D. | 0.997 | 1.000 | 0.443 | 455 |
| MMseqs2 | Cov. mode 1 | Elec. | 1.000 | 0.999 | 0.637 | 178 |
| | | M.D. | 1.000 | 1.000 | 0.999 | 174 |
| | Default | Elec. | 1.000 | 0.999 | 0.584 | 196 |
| | | M.D. | 1.000 | 1.000 | 0.982 | 162 |
| | 50% minimum identity | Elec. | 1.000 | 0.999 | 0.584 | 184 |
| | | M.D. | 1.000 | 1.000 | 0.982 | 174 |
| Starcode | Default | Elec. | 1.000 | 0.997 | 0.076 | 489 |
| | | M.D. | 1.000 | 1.000 | 0.450 | 133 |
| | Sphere clustering | Elec. | 1.000 | 0.999 | 0.227 | 661 |
| | | M.D. | 1.000 | 1.000 | 0.895 | 149 |
| | Sphere clustering<br>Distance 6 | Elec. | 1.000 | 0.999 | 0.276 | 1710 |
| | | M.D. | 1.000 | 1.000 | 0.903 | 194 |

**Supplementary Table 2: Overview of codecs for DNA data storage in the literature.**[7,8] Note that the cut-off date for consideration was October 2023.

| Selected? | Name | Year | Inner EC | Outer EC | Constraints | In-vitro? | Code? | Ref. | Comments |
|---|---|---|---|---|---|---|---|---|---|
| | Church et al. | 2012 | None | None | HP | Yes | Yes | 9 | No error-correction component |
| Yes | Goldman et al. | 2013 | Parity | Repetition | GC, HP | Yes | Yes | 10 | |
| | Grass et al. | 2015 | RS | RS | GC, HP | Yes | | 11 | Superseded by DNA-RS |
| | Yazdi et al. | 2015 | None | None | GC, ΔG | Yes | | 12 | Uses ultra-long sequences |
| | Bornholt et al. | 2016 | Parity | XOR/Repetition | None | Yes | | 13 | No implementation available |
| | Blawat et al. | 2016 | BCH | RS | GC, HP | Yes | | 14 | No implementation available |
| | Yazdi et al. | 2017 | BCH | None | GC | Yes | Yes | 15 | Uses ultra-long sequences |
| Yes | DNA Fountain | 2017 | RS | Fountain | GC, HP | Yes | Yes | 16 | |
| | Organick et al. | 2018 | None | RS | HP | Yes | | 17 | No implementation available |
| | Oligoarchive | 2019 | Parity | Repetition | GC, HP | Yes | | 18 | Specific to database structures |
| | RA code | 2019 | CRC | RA | GC, HP | Yes | | 19 | No implementation available |
| | Large LDPC | 2019 | LDPC | | None | Yes | Yes | 20 | Employs single block code |
| | Deng et al. | 2019 | LDPC | None | GC, HP | | | 21 | No implementation available |
| | Wang et al. | 2019 | None | None | GC, HP | | | 22 | No implementation available |
| | Anavy et al. | 2019 | RS | Fountain | GC, HP | Yes | Yes | 23 | Uses degenerate sequences |
| | Choi et al. | 2019 | None | RS | HP | Yes | | 24 | Uses degenerate sequences |
| Yes | HEDGES | 2020 | HEDGES | RS | GC, HP | Yes | Yes | 25 | |
| Yes | DNA-RS | 2020 | RS | RS | None | Yes | Yes | 26,27 | |
| | Lenz et al. | 2020 | multiple | LDPC | None | | | 28 | No implementation available |
| | JPEG | 2021 | Parity | None | GC, HP | Yes | | 29 | Specific to image storage |
| | Chen et al. | 2021 | LDPC | None | GC | Yes | | 30 | Designed for artificial chromosome |
| Yes | Yin-Yang | 2022 | None | None | GC, HP, ΔG | Yes | Yes | 31 | |
| | DBGPS | 2022 | CRC | Fountain | HP, kmer | Yes | Yes | 32 | Intended for strand reconstruction |
| | 2DDNA | 2022 | None | LDPC | GC, HP | Yes | Yes | 33 | Uses backbone for encoding |
| Yes | DNA-Aeon | 2023 | AC-based | Fountain | GC, HP, motifs | Yes | Yes | 34 | |
| | MAFFT | 2023 | MSA | None | None | | Yes | 35 | Uses only multiple-sequence alignment |
| | Zan et al. | 2023 | MSA | None | GC, HP | | Yes | 36 | No in-vitro experiment |
| | Zhao et al. | 2024 | HEDGES | RS | GC, HP | Yes | Yes | 37 | Uses degenerate sequences, for nanopores |

**Supplementary Table 3: Full results of all codec-clustering combinations.** The best-performing parameter set of each clustering algorithm was paired with each codec in the basic error scenario, yielding the error rate at which decoding succeeded after clustering with 95% probability. For each codec and code rate, the best-performing clustering algorithm was selected for all further studies, and is indicated by green shading.

| Codec | Code rate | Naive | CD-Hit | Clover | LSH | Starcode |
|---|---|---|---|---|---|---|
| | bit nt$^{-1}$ | Default | 85% identity | D15V4 | Default | Sphere, distance 6 |
| DNA-Aeon | 0.50 | 0.002 | 0.066 | 0.029 | 0.024 | 0.014 |
| | 1.00 | 0.003 | 0.074 | 0.042 | 0.035 | 0.015 |
| | 1.50 | 0.002 | 0.077 | 0.042 | 0.037 | 0.017 |
| DNA Fountain | 0.50 | 0.017 | 0.032 | - | - | 0.045 |
| | 1.00 | 0.016 | 0.027 | - | - | 0.042 |
| | 1.50 | 0.010 | 0.051 | - | - | - |
| DNA-RS | 0.50 | 0.033 | 0.119 | 0.107 | 0.107 | 0.046 |
| | 1.00 | 0.022 | 0.119 | 0.091 | 0.107 | 0.048 |
| | 1.50 | 0.017 | 0.103 | 0.086 | 0.091 | 0.043 |
| Goldman | 0.34 | 0.016 | - | - | 0.067 | 0.023 |
| HEDGES | 0.63 | 0.077 | 0.109 | - | 0.133 | 0.086 |
| | 1.07 | 0.022 | 0.120 | - | 0.103 | 0.049 |
| Yin-Yang | 1.85 | - | 0.042 | - | - | - |

**Supplementary Table 4: Selected codec parameters for codec "DNA-Aeon".**

| Parameter | Default | In-silico studies | | | In-vitro pool experiment | | |
|---|---|---|---|---|---|---|---|
| | | *Low* | *Medium* | *High* | *Medium* | *High* | *Max* |
| Homopolymer | 4 | 4 | 4 | 4 | 4 | 4 | 4 |
| GC-content | 0.4-0.6 | 0.0-1.0 | 0.0-1.0 | 0.0-1.0 | 0.0-1.0 | 0.0-1.0 | 0.0-1.0 |
| Package redundancy | 0.45 | 1.68 | 0.34 | 0.031 | 0.32 | 0.028 | 0.0 |
| Chunk size | 14 | 25 | 25 | 28 | 20 | 24 | 28 |
| Sync value | 4 | 4 | 4 | 8 | 4 | 12 | 0 |
| Error correction | CRC | CRC | CRC | CRC | CRC | CRC | nocode |
| Codeword length | 10 | 10 | 10 | 10 | 10 | 10 | 10 |
| CRC threshold | 3 | 3 | 3 | 3 | 3 | 3 | 3 |
| Loop | 1 | 1 | 1 | 1 | 1 | 1 | 1 |
| Finish | 0 | 0 | 0 | 0 | 0 | 0 | 0 |
| Penalty (CRC) | 0.1 | 0.1 | 0.1 | 0.1 | 0.1 | 0.1 | 0.1 |
| Penalty (No-Hit) | 8 | 8 | 8 | 8 | 8 | 8 | 8 |

**Supplementary Table 5: Selected codec parameters for codec "DNA Fountain".**

| Parameter | Default | In-silico studies | | | In-vitro pool experiment | | |
|---|---|---|---|---|---|---|---|
| | | *Low* | *Medium* | *High* | *Medium* | *High* | *Max* |
| Alpha | 0.07 | 2.35 | 0.68 | 0.19 | 0.6 | 0.14 | 0.0 |
| Payload | 32 | 32 | 32 | 34 | 24 | 26 | 27 |
| RS length | - | 2 | 2 | 0 | 2 | 1 | 0 |
| Hamming distance | 100 | 100 | 100 | 100 | 100 | 100 | 100 |
| GC-content | - | 0.0-1.0 | 0.0-1.0 | 0.0-1.0 | 0.0-1.0 | 0.0-1.0 | 0.0-1.0 |
| Homopolymer | 4 | 4 | 4 | 4 | 4 | 4 | 4 |
| Delta | 0.05 | 0.1 | 0.1 | 0.1 | 0.1 | 0.1 | 0.1 |
| C-Dist | 0.1 | 0.025 | 0.025 | 0.025 | 0.025 | 0.025 | 0.025 |
| Header size | 4 | 4 | 4 | 4 | 4 | 4 | 4 |

**Supplementary Table 6: Selected codec parameters for codec "DNA-RS".** As the number of sequences is dependent on the file size $s$, it is parameterized according to the equations below the table.

| Parameter | Default | In-silico studies | | | In-vitro pool experiment | | |
|---|---|---|---|---|---|---|---|
| | | *Low* | *Medium* | *High* | *Medium* | *High* | *Max* |
| Mi | 6 | 6 | 6 | 6 | 6 | 6 | 6 |
| Mo | 14 | 14 | 14 | 14 | 14 | 14 | 14 |
| Index | 24 | 24 | 24 | 24 | 24 | 24 | 24 |
| Number of seqs. | - | $f_1(s)$ | $f_2(s)$ | $f_3(s)$ | 1110 | 823 | 928 |
| Sequence length | - | 150 | 144 | 144 | 126 | 126 | 102 |
| Inner red. symbols | - | 4 | 2 | 2 | 3 | 3 | 2 |

$$f_1(s) = \left\lfloor \frac{875}{8192} s \right\rfloor, \qquad f_2(s) = \left\lfloor \frac{855}{15360} s \right\rfloor, \qquad f_3(s) = \left\lfloor \frac{570}{15360} s \right\rfloor$$

**Supplementary Table 7: Selected codec parameters for codec "HEDGES".** As the sequence length is dependent on the file size $s$ and the number of packets, it is parameterized according to the equations below the table.

| Parameter | Default | In-silico studies | | In-vitro pool experiment |
|---|---|---|---|---|
| | | *Low* | *Medium* | |
| Code rate index | - | 3 | 1 | 1 |
| Sequence length | - | $f_1(s)$ | $f_2(s)$ | 110 |
| Homopolymer | 4 | 4 | 4 | 4 |
| GC window | 4-8 | 4-8 | 4-8 | 4-8 |

$$f_1(s) = 28.0\, s - 1015, \qquad f_2(s) = 44.6\, s - 1338$$

**Supplementary Table 8: Selected codec parameters for codec "Yin-Yang".**

| Parameter | Default | In-silico studies | In-vitro pool experiment |
|---|---|---|---|
| Homopolymer | 4 | 4 | 4 |
| GC-content | 0.6 | 0.75 | 0.75 |
| Search count | 100 | 100 | 100 |
| Segment length | 120 | 140 | 110 |

**Supplementary Table 9: Comparisons of achieved storage densities.** Note that both Organick et al.[2] and Grass et al.[11,38] report slightly different storage densities in their respective studies (i.e., 17 EB $g^{-1}$ by Organick et al.[2]). This is due to different assumptions for calculation, see Supplementary Note 2. In this table, all calculations were harmonized to facilitate fair comparisons. Synthesis providers are abbreviated to TW (Twist Biosciences) and GS (Genscript).

| **Codec** | **DNA-Aeon** | | **DNA-RS** | | **Literature** | |
|---|---|---|---|---|---|---|
| **Scenario** | **High-F.** | **Low-F.** | **High-F.** | **Low-F.** | **Organick et al.**[2] | **Grass et al.**[11,38] |
| File size / bit | 139 264 | | 139 264 | | 255 512[a] | 663 168 |
| Sequence count | 1 150 | | 1 110 | | 2 042 | 4 991 |
| Min. phys. red. | 2.0 | 6.6 | 2.0 | 6.6 | 10 | 3898 |
| Sequencing depth | 30 | | 30 | | 35 | 372 |
| State | double-stranded | | double-stranded | | single-stranded | double-stranded |
| Synthesis provider | TW | GS | TW | GS | TW | GS |
| ***Considering only payload*** | | | | | | |
| Length / nt | 120 | | 126 | | 110 | 117 |
| Code rate / bit $nt^{-1}$ | 1.01 | | 1.00 | | 1.14 | 1.14 |
| Stor. dens. / EB $g^{-1}$ | 57.4 | 17.4 | 56.9 | 17.2 | 25.9 | 0.033 |
| ***Considering payload and primer adapters*** | | | | | | |
| Length / nt | 161 | | 167 | | 150 | 158 |
| Code rate / bit $nt^{-1}$ | 0.75 | | 0.76 | | 0.83 | 0.84 |
| Stor. dens. / EB $g^{-1}$ | 42.6 | 13.0 | 43.2 | 12.9 | 19.0 | 0.025 |
| ***Considering all nucleotides, including suffix and padding*** | | | | | | |
| Length / nt | 170 | | 170 | | 150 | 158 |
| Code rate / bit $nt^{-1}$ | 0.71 | | 0.74 | | 0.83 | 0.84 |
| Stor. dens. / EB $g^{-1}$ | 40.4 | 12.3 | 42.1 | 12.7 | 19.0 | 0.025 |

[a] The exact file size was extracted from the SI in Ref. [17] using the random-access primers described by both studies, matching file 10. The file size of 0.1 KB reported in the original study[2] does not match the reported storage density and code rate, and was thus assumed to be erroneous.

**Supplementary Table 10: Sequence properties per codec and code rate for the pool experiment.**

| Codec | File size | Code rate | Count | Ratio | Length | Suffix |
|---|---|---|---|---|---|---|
| | kB | bit nt$^{-1}$ | # seqs. | % | nt | |
| DNA-Aeon | 17 | 1.01 | 1150 | 10.2 | 120 | AGG |
| | 19 | 1.51 | 834 | 7.4 | 124 | ACC |
| | 19 | 1.81 | 695 | 6.2 | 124 | AAA |
| DNA Fountain | 17 | 1.00 | 1162 | 10.3 | 120 | CCA |
| | 19 | 1.47 | 854 | 7.6 | 124 | CAC |
| | 19 | 1.74 | 722 | 6.4 | 124 | ATT |
| DNA-RS | 17 | 1.00 | 1110 | 9.8 | 126 | GAG |
| | 19 | 1.50 | 823 | 7.3 | 126 | CTG |
| | 19 | 1.64 | 928 | 8.2 | 102 | CGT |
| Goldman | 5 | 0.34 | 1032 | 9.1 | 117 | TAT |
| HEDGES | 17 | 0.99 | 1275 | 11.3 | 110 | GTC |
| Yin-Yang | 19 | 1.82 | 708 | 6.3 | 121 | TCG |

**Supplementary Table 11: Primer sequences used for amplification, qPCR, and sequencing preparation in this study.**

| Name | Purpose | Sequence |
|---|---|---|
| 0F | Amplification, qPCR | ACACGACGCTCTTCCGATCT |
| 0R | Amplification, qPCR | AGACGTGTGCTCTTCCGATCT |
| 2FUF | Sequencing prep. | AATGATACGGCGACCACCGAGATCTACACTCTTTCCCTACACGACGCTCTTCCGATCT |
| 2RIF-GM5 | Sequencing prep. | CAAGCAGAAGACGGCATACGAGATCACTGTGTGACTGGAGTTCAGACGTGTGCTCTTCCGATCT |
| 2RIF-GM6 | Sequencing prep. | CAAGCAGAAGACGGCATACGAGATATTGGCGTGACTGGAGTTCAGACGTGTGCTCTTCCGATCT |
| 2RIF-GM7 | Sequencing prep. | CAAGCAGAAGACGGCATACGAGATGATCTGGTGACTGGAGTTCAGACGTGTGCTCTTCCGATCT |
| 2RIF-GM8 | Sequencing prep. | CAAGCAGAAGACGGCATACGAGATTCAAGTGTGACTGGAGTTCAGACGTGTGCTCTTCCGATCT |
| 2RIF-GM11 | Sequencing prep. | CAAGCAGAAGACGGCATACGAGATGTAGCCGTGACTGGAGTTCAGACGTGTGCTCTTCCGATCT |

**Supplementary Table 12: qPCR results of experiments in the worst-case scenario.** Coverage is calculated based on the calibration curve obtained by serial dilution, shown in Supplementary Figure 12. A standard from the calibration curve was used to standardize the experimentally measured cycle threshold. Each sample was measured in duplicate.

| Sample | qPCR Cq | qPCR $Cq_{std}$ | Calc. coverage | Mean coverage |
|---|---:|---:|---:|---:|
| Standard | 17.26 | 17.435 | 128 | 128 |
| | 17.28 | 17.455 | 127 | |
| Cov. 5 | 21.85 | 22.025 | 6.60 | 6.56 |
| | 21.87 | 22.045 | 6.51 | |
| Cov. 10 | 20.85 | 21.025 | 12.6 | 12.5 |
| | 20.88 | 21.055 | 12.4 | |
| Cov. 25 | 19.67 | 19.845 | 27.1 | 26.9 |
| | 19.69 | 19.865 | 26.7 | |
| Cov. 50 | 18.62 | 18.795 | 53.4 | 52.0 |
| | 18.70 | 18.875 | 50.7 | |
| Cov. 1000 | 14.01 | 14.185 | 1054 | 1054 |
| | 14.01 | 14.185 | 1054 | |

**Supplementary Table 13: qPCR results of experiments in the best-case scenario.** Coverage is calculated based on the calibration curve obtained by serial dilution, shown in Supplementary Figure 12. A standard from the calibration curve was used to standardize the experimentally measured cycle threshold. Each sample was measured in duplicate.

| Sample | qPCR Cq | qPCR $Cq_{std}$ | Calc. coverage | Mean coverage |
|---|---:|---:|---:|---:|
| Standard | 17.09 | 16.785 | 137 | 137 |
| | 17.09 | 16.785 | 137 | |
| Cov. 2 | 23.54 | 23.235 | 1.99 | 1.96 |
| | 23.58 | 23.275 | 1.94 | |
| Cov. 5 | 22.16 | 21.855 | 4.91 | 4.93 |
| | 22.15 | 21.845 | 4.95 | |
| Cov. 10 | 21.20 | 20.895 | 9.23 | 9.23 |
| | 21.20 | 20.895 | 9.23 | |
| Cov. 25 | 19.33 | 19.025 | 31.5 | 31.2 |
| | 19.36 | 19.055 | 30.9 | |
| Cov. 1000 | 13.70 | 13.395 | 1265 | 1273 |
| | 13.68 | 13.375 | 1281 | |

### Supplementary Figures

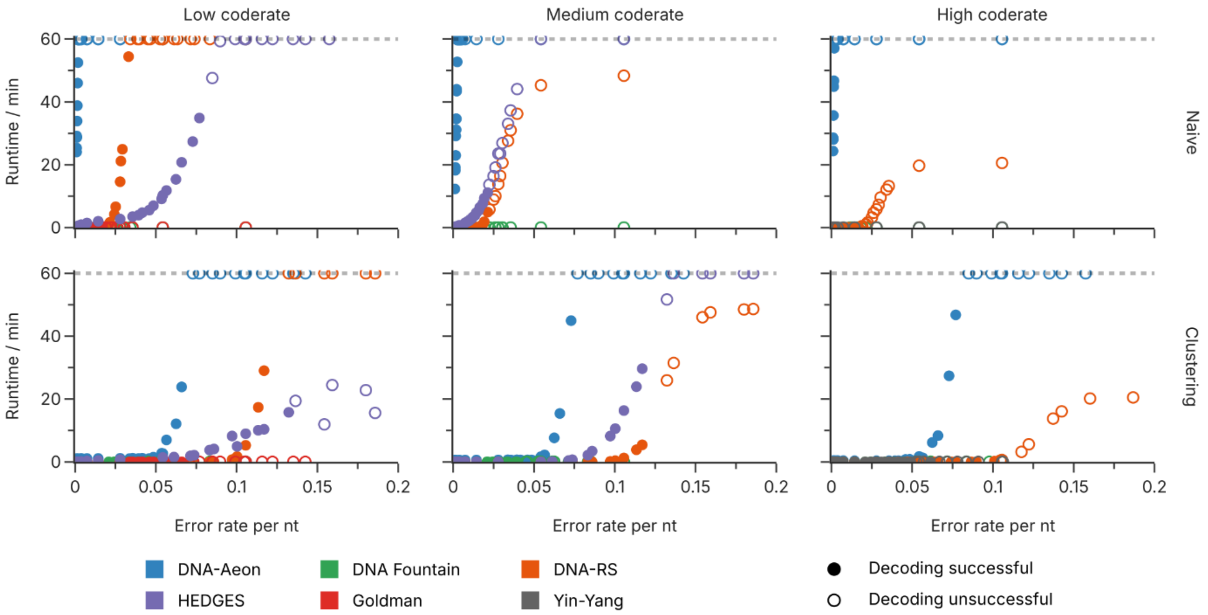


**Supplementary Figure 1: Decoding time as a function of error rate in the basic error scenario with combined errors.** The runtime of the decoding step is shown for the DNA-Aeon, DNA Fountain, DNA-RS, Goldman, HEDGES, and Yin-Yang codecs at all used code rates, when substitutions, deletions, and insertions are introduced simultaneously at a ratio of 53:45:2. Points correspond to individual runs of the pipeline. Open circles denote individual runs which failed the decoding step, either due to violation of the runtime constraint or due to insufficient error-correction capabilities.

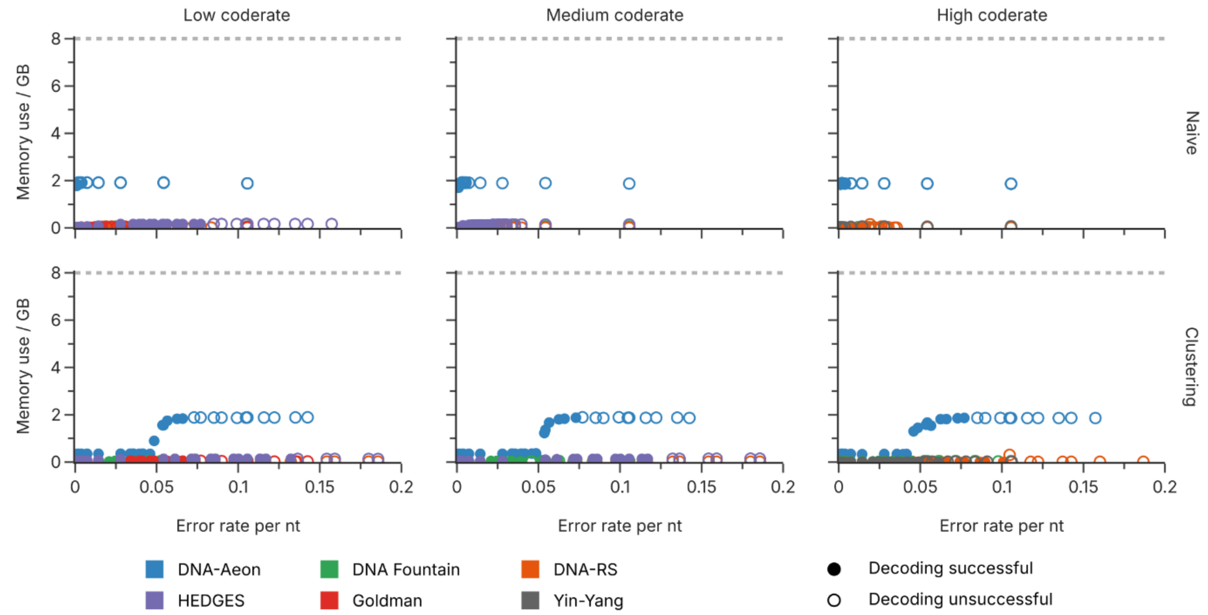


**Supplementary Figure 2: Memory use as a function of error rate in the basic error scenario with combined errors.** The memory use of the decoding step is shown for the DNA-Aeon, DNA Fountain, DNA-RS, Goldman, HEDGES, and Yin-Yang codecs at all used code rates, when substitutions, deletions, and insertions are introduced simultaneously at a ratio of 53:45:2. Points correspond to individual runs of the pipeline. Open circles denote individual runs which failed the decoding step, either due to violation of the runtime constraint or due to insufficient error-correction capabilities.

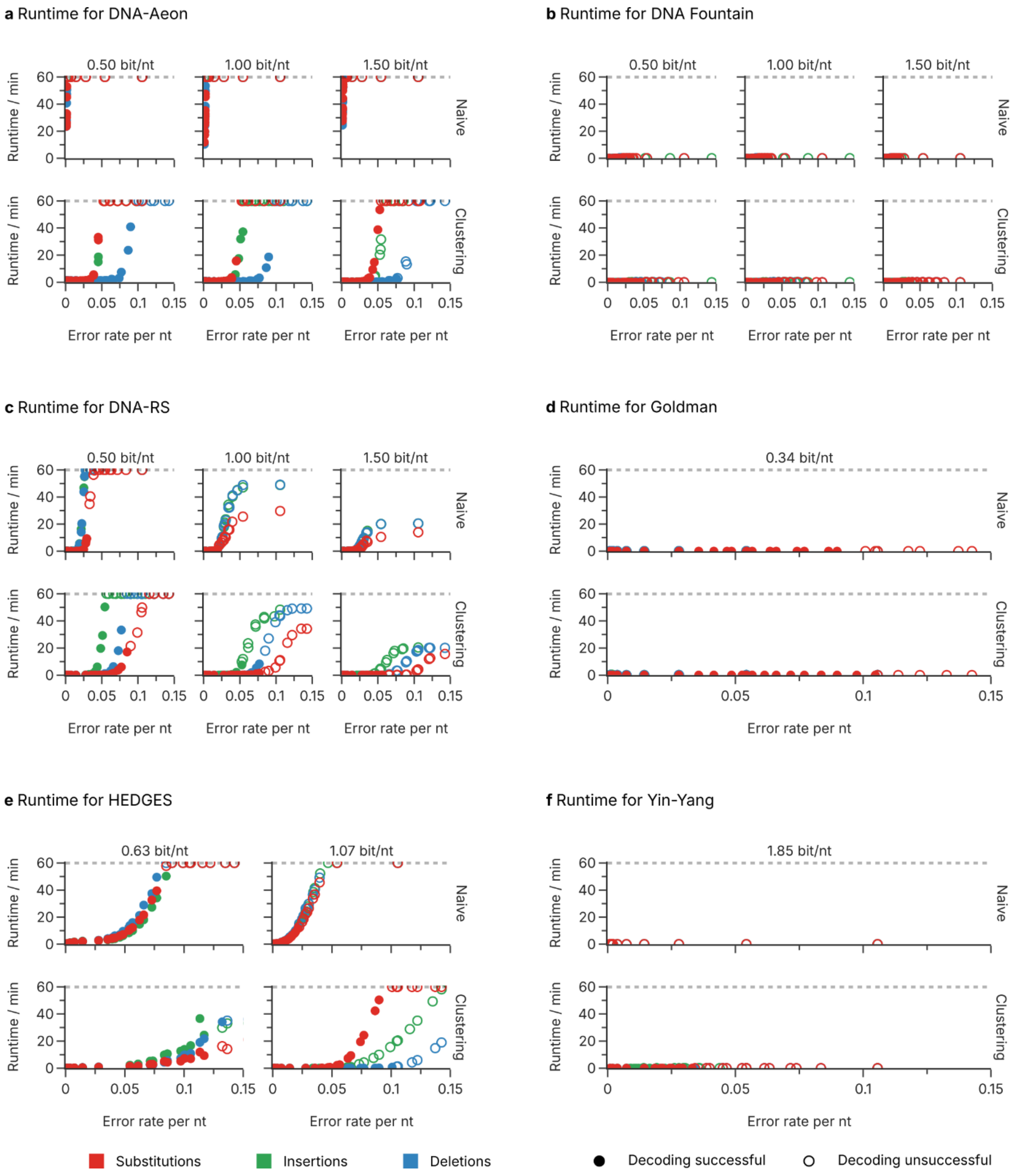


**Supplementary Figure 3: Decoding time as a function of error rate in the basic error scenario with individual errors.** The runtime of the decoding step is shown for the DNA-Aeon (**a**), DNA Fountain (**b**), DNA-RS (**c**), Goldman (**d**), HEDGES (**e**), and Yin-Yang (**f**) codecs at all used code rates, when substitutions (red), deletions (blue), or insertions (green) are introduced individually. Points correspond to individual runs of the pipeline at the specified error rate and error type. Open circles denote individual runs which failed the decoding step, either due to violation of the runtime constraint or due to insufficient error-correction capabilities.

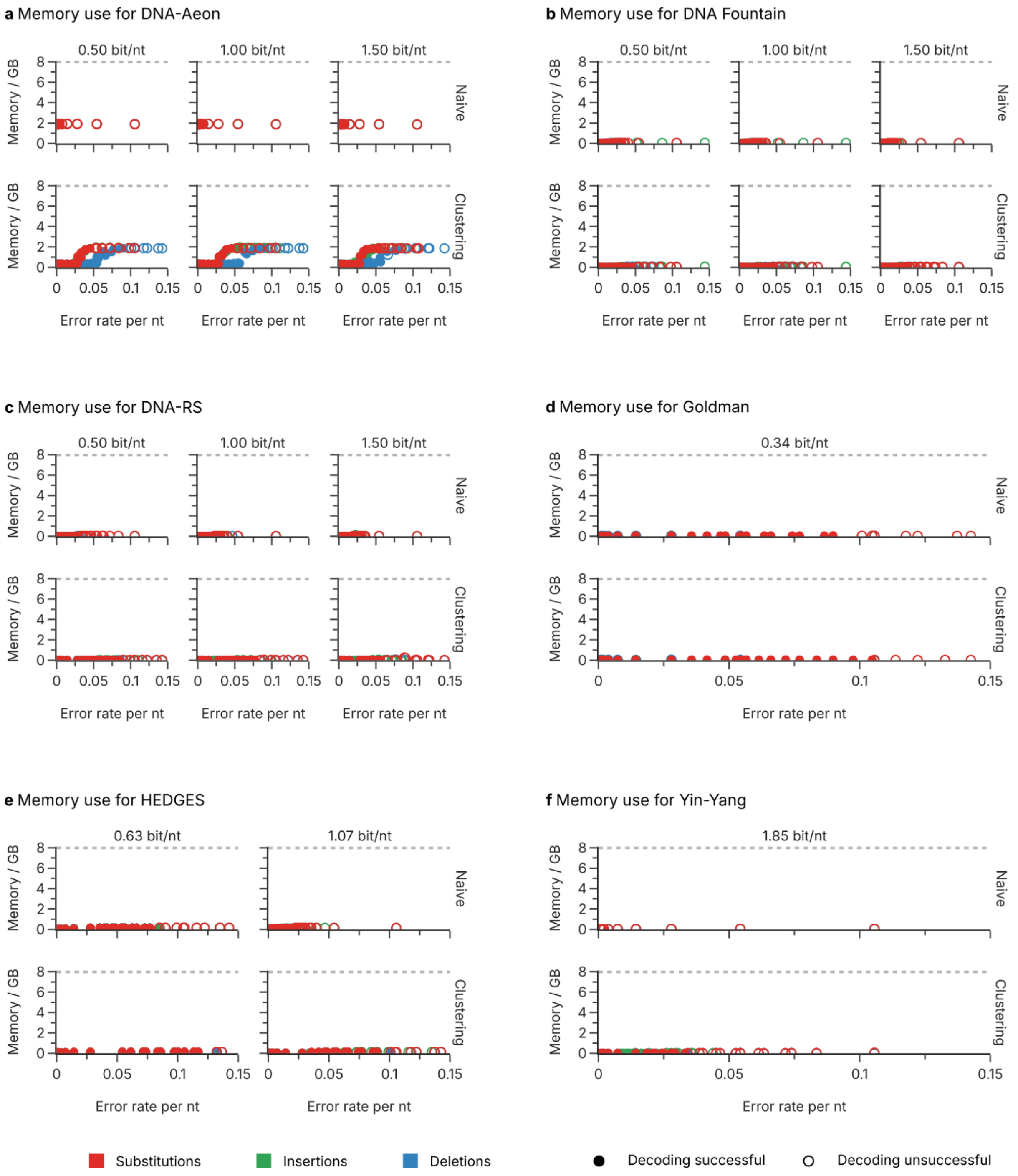


**Supplementary Figure 4: Memory use as a function of error rate in the basic error scenario with individual errors.** The memory use of the decoding step is shown for the DNA-Aeon (**a**), DNA Fountain (**b**), DNA-RS (**c**), Goldman (**d**), HEDGES (**e**), and Yin-Yang (**f**) codecs at all used code rates, when substitutions (red), deletions (blue), or insertions (green) are introduced individually. Points correspond to individual runs of the pipeline at the specified error rate and error type. Open circles denote individual runs which failed the decoding step, either due to violation of the runtime constraint or due to insufficient error-correction capabilities.

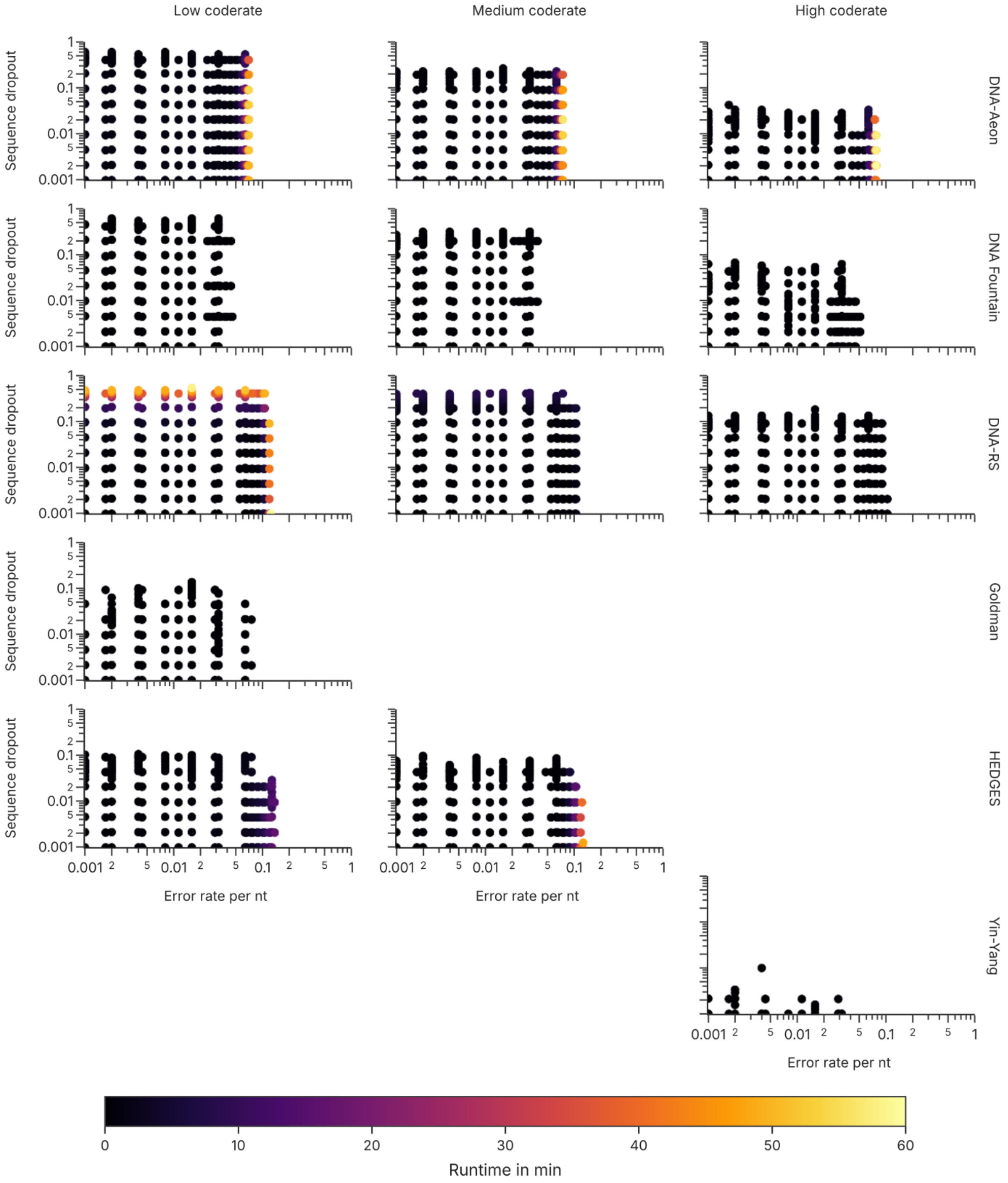


**Supplementary Figure 5: Decoding time as a function of overall error rate and sequence dropout in the two-parameter sensitivity analysis.** The runtime of the decoding step is shown for the DNA-Aeon, DNA Fountain, DNA-RS, Goldman, HEDGES, and Yin-Yang codecs (top to bottom), at the different code rates (left to right). Points correspond to individual runs of the pipeline at the specified error rate and sequence dropout. Only individual runs which led to successful decoding are shown (i.e., runs which violated the runtime constraint or failed due to insufficient error-correction capabilities are not shown).

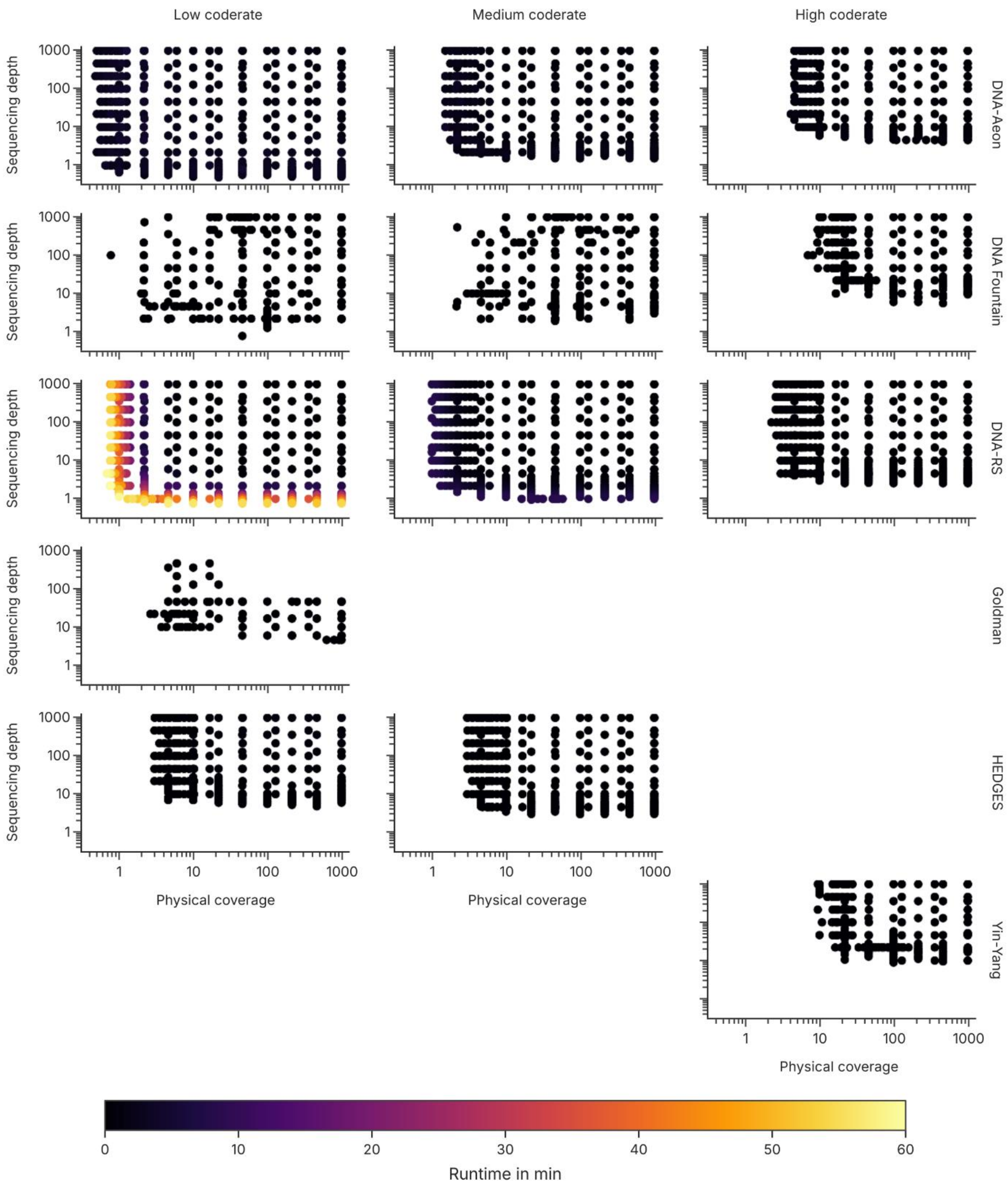


**Supplementary Figure 6: Decoding time as a function of sequencing depth and physical coverage in the high-fidelity scenario.** The runtime of the decoding step is shown for the DNA-Aeon, DNA Fountain, DNA-RS, Goldman, HEDGES, and Yin-Yang codecs (top to bottom), at the different code rates (left to right). Points correspond to individual runs of the pipeline at the specified physical coverage and sequencing depth of the best-case scenario. Only individual runs which led to successful decoding are shown (i.e., runs which violated the runtime constraint or failed due to insufficient error-correction capabilities are not shown).

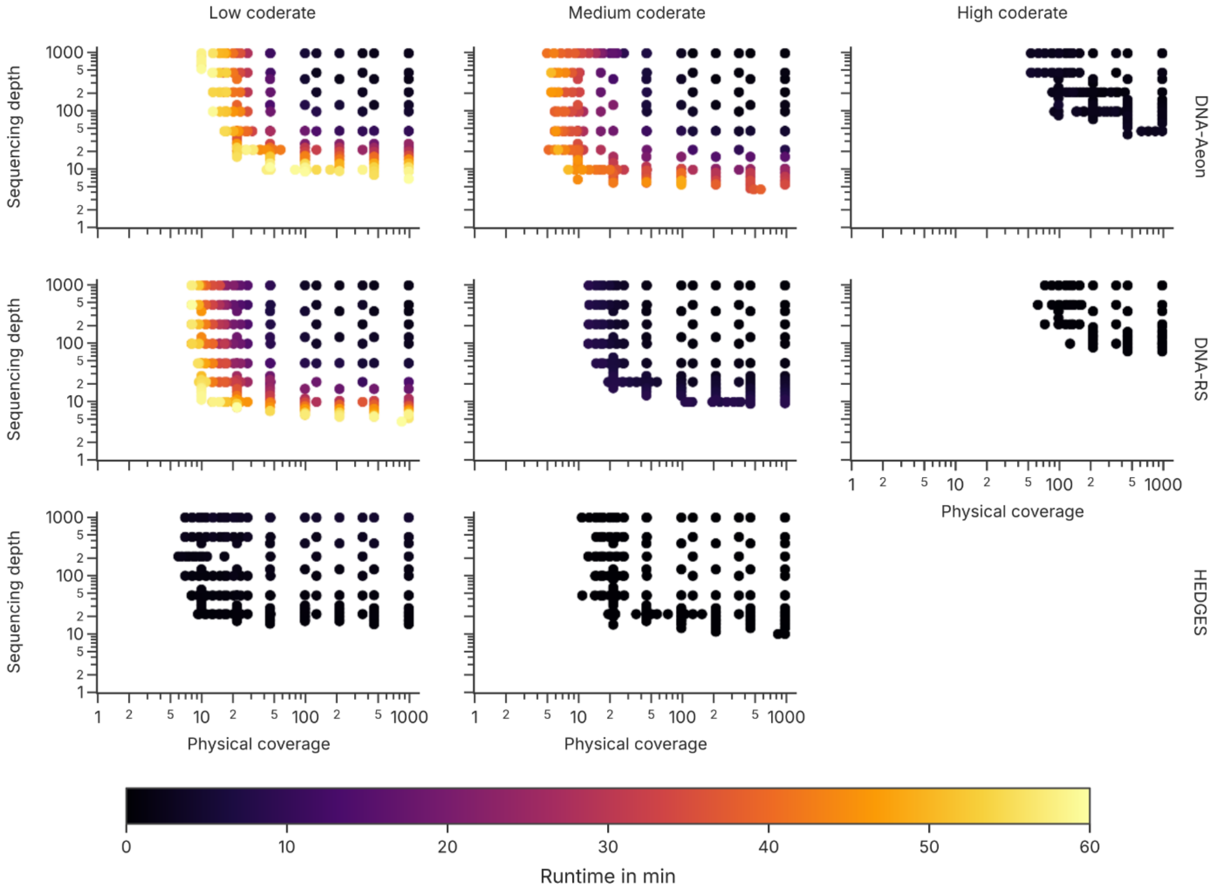


**Supplementary Figure 7: Decoding time as a function of sequencing depth and physical coverage in the low-fidelity scenario.** The runtime of the decoding step is shown for the DNA-Aeon, DNA-RS, and HEDGES codecs (top to bottom, other codecs failed to successfully decode at all), at the different code rates (left to right). Points correspond to individual runs of the pipeline at the specified physical coverage and sequencing depth of the worst-case scenario. Only individual runs which led to successful decoding are shown (i.e., runs which violated the runtime constraint or failed due to insufficient error-correction capabilities are not shown).

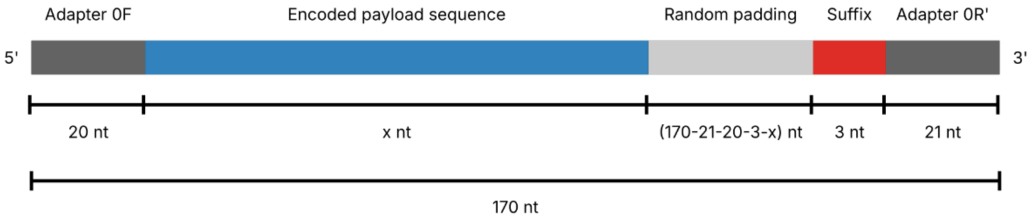


**Supplementary Figure 8: Sequence design for pool experiments.** The sequences are composed of the forward and reverse adapter for amplification (dark gray), the encoded payload sequence as generated by a codec (blue), a random padding to pad the combined sequence to 170 nt (light gray), and a short suffix for identification (red). See Supplementary Table 10 or additional information on sequence properties of the individual codecs.

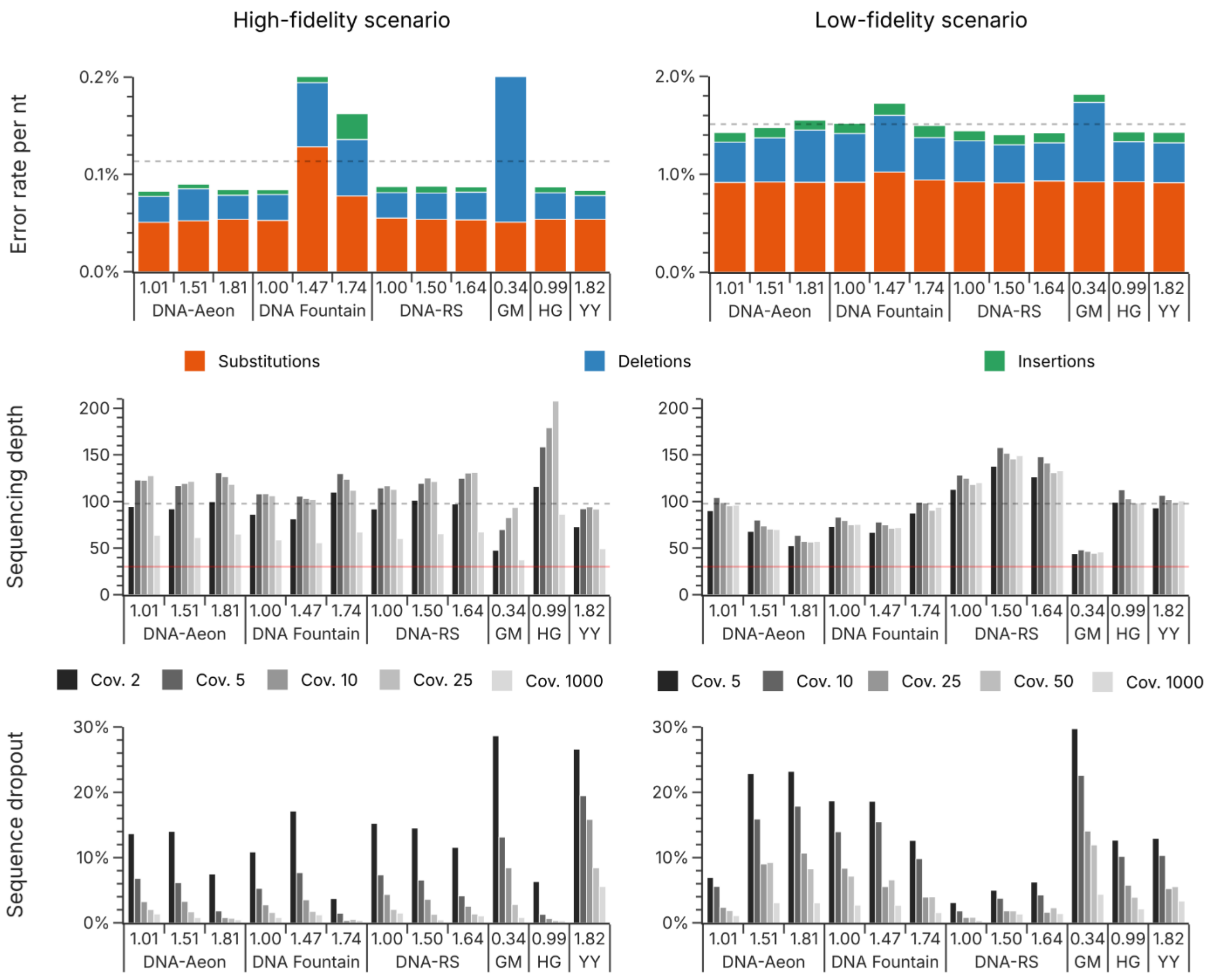


**Supplementary Figure 9: Comparison of the mean error rates, sequencing depths, and rates of sequence dropout between codecs and scenarios in the pool experiments.** The error rates (top row) in the high- (left) and low-fidelity scenario (right) are composed of substitutions (orange), deletions (blue), and insertions (green). In each case, the sequencing data from the experiment with a coverage of 1000x was used for error analysis. The dotted line represents the mean across all codecs. The sequencing depth highlights differences between codecs (groups) and coverages (individual colors). The mean sequencing depth (middle row) across all codecs is shown with a dotted line. The solid red line indicates a sequencing depth of 30x, to which all sequencing data was downsampled for the decoding experiments, as well as the analysis of sequence dropout. The sequence dropout (bottom row) shows differences between codecs (groups) and coverages (individual colors). To quantify the sequence dropout, the full sequencing data was downsampled to a sequencing depth of 30x ten times, and the average fraction of sequences without a corresponding read are reported.

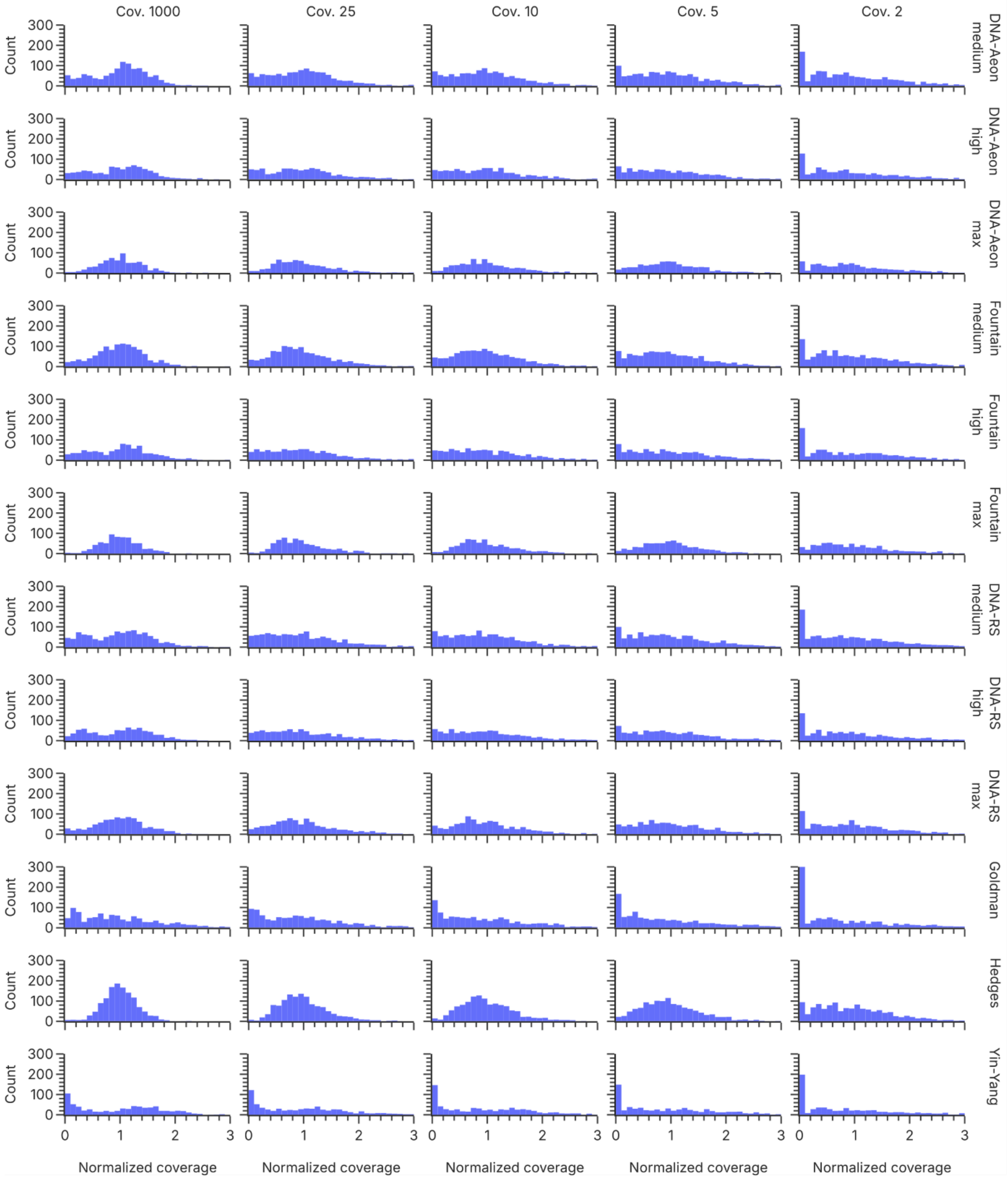


**Supplementary Figure 10: Coverage distributions between codecs and coverages in the pool experiment using the high-fidelity scenario.** The histograms show the homogeneity of the sequence coverage, normalized to the mean coverage. The more skewed the coverage distribution, the less homogeneous the representation of each sequence in the oligo pool.

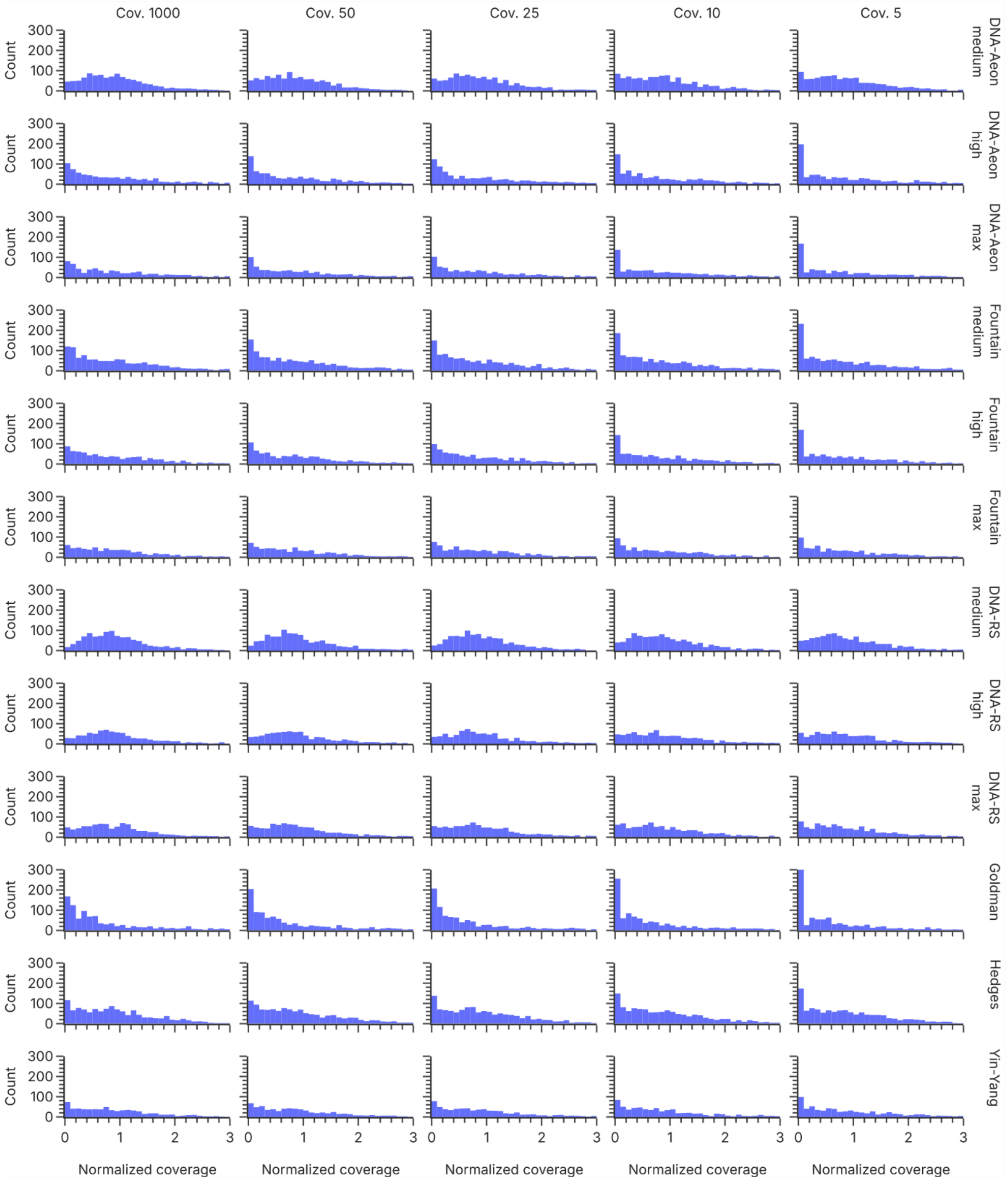


**Supplementary Figure 11: Coverage distributions between codecs and coverages in the pool experiment using the low-fidelity scenario.** The histograms show the homogeneity of the sequence coverage, normalized to the mean coverage. The more skewed the coverage distribution, the less homogeneous the representation of each sequence in the oligo pool.

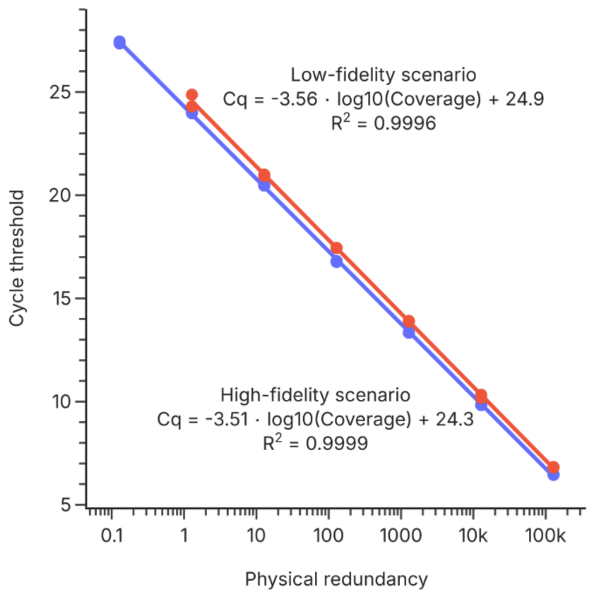


**Supplementary Figure 12: qPCR calibration curves in the high- and low-fidelity scenarios.** The calibration curves were generated by serial dilutions of the two master pools generated from the first amplification PCR after synthesis. Conversion of mass concentration to coverage was performed assuming a physical coverage of 509074x per ng, and a sample volume of 5 µL (see Methods).
